# An insulin-stimulated proteolytic mechanism links energy expenditure with glucose uptake

**DOI:** 10.1101/817049

**Authors:** Estifanos N. Habtemichael, Don T. Li, João Paulo Camporez, Xavier O. Westergaard, Chloe I. Sales, Xinran Liu, Francesc López-Giráldez, Stephen G. DeVries, Hanbing Li, Diana M. Ruiz, Kenny Y. Wang, Bhavesh S. Sayal, Sandro Hirabara, Daniel F. Vatner, William Philbrick, Gerald I. Shulman, Jonathan S. Bogan

## Abstract

Mechanisms to coordinately regulate energy expenditure and glucose uptake into muscle and fat cells are not well described. Insulin stimulates glucose uptake in part by causing site-specific endoproteolytic cleavage of TUG, which mobilizes GLUT4 glucose transporters to the cell surface. Here, we show that the TUG C-terminal cleavage product enters the nucleus, binds the transcriptional regulators PGC-1α and PPARγ, and increases oxidative metabolism and thermogenic protein expression. Muscle-specific genetic manipulation of this pathway impacts whole-body energy expenditure, independent of glucose uptake. The PPARγ2 Pro12Ala polymorphism, which reduces diabetes risk, enhances TUG binding. The TUG cleavage product stabilizes PGC-1α and is itself susceptible to an Ate1 arginyltransferase -dependent degradation mechanism; binding of the TUG product confers Ate1-dependent stability upon PGC-1α. We conclude that TUG cleavage coordinates energy expenditure with glucose uptake, that this pathway may contribute to the thermic effect of food, and that its attenuation may be important in obesity.

## Introduction

How energy expenditure in mammals is linked to the availability of metabolic substrates is not well understood. Although it is a primary nutritional signal, insulin has not been implicated in direct control of thermogenic mechanisms. Insulin stimulates glucose uptake into muscle and fat cells by mobilizing GLUT4 glucose transporters to the plasma membrane (Bogan, 2012; Leto and Saltiel, 2012). In unstimulated cells, GLUT4 is trapped intracellularly by TUG proteins (Bogan et al., 2003; Yu et al., 2007). Insulin liberates this pool of “GLUT4 storage vesicles” (GSVs) by triggering site-specific endoproteolytic cleavage of TUG (Belman et al., 2014; Bogan et al., 2012; Löffler et al., 2013). In adipocytes, this process requires the Usp25m protease and is impaired in rodents with diet-induced insulin resistance (Habtemichael et al., 2018).

Impaired regulation of GSVs by TUG may contribute to multiple aspects of the metabolic syndrome. In addition to GLUT4, TUG-regulated GSVs contain the aminopeptidase IRAP, which inactivates circulating vasopressin, and lipoprotein receptors (LRP1, sortilin), which may act in cholesterol metabolism (Habtemichael et al., 2015; Li et al., 2019). Transgenic mice with constitutive, insulin-independent TUG cleavage in muscle exhibit not only increased fasting glucose uptake, due to GLUT4, but also increased water intake, due to IRAP. These mTUG^UBX-Cter^ mice (here called “UBX mice”) also display increased energy expenditure (Löffler et al., 2013), which remains unexplained. Here, we report that the TUG C-terminal cleavage product mediates this effect by promoting the expression of genes that enhance oxidative metabolism.

## Results

In UBX mice, muscle-specific transgenic expression of an unstable TUG fragment recruits PIST, a negative regulator of TUG cleavage, for proteosomal degradation, and endogenous intact TUG proteins are cleaved constitutively in the absence of an insulin signal (Löffler et al., 2013). In fasting UBX mice, GSVs are translocated and TUG cleavage products are generated in muscle. Thus, the 12-13% increase in energy expenditure observed in these mice could result from translocation of a GSV cargo protein or from effects of a cleavage product. To distinguish these possibilities, we created muscle-specific TUG knockout (MTKO) mice. We predicted that in the absence of intact TUG, GSV cargoes would be translocated to the cell surface in muscle cells, as in adipocytes (Yu et al., 2007), but no TUG cleavage products will be generated. With respect to production of the TUG C-terminal cleavage product, UBX and MTKO mice are gain- and loss-of-function models, respectively.

TUG deletion in muscle had no effect on GLUT4 or IRAP abundances, or on body weight and composition (Figures 1A, 1B, and S1A–S1C). Like UBX mice (Löffler et al., 2013), MTKO mice had reduced fasting plasma glucose and insulin concentrations, compared to wildtype (WT) controls (Figures 1C and 1D). Dynamic measurements of glucose flux showed that during fasting, whole-body glucose turnover was increased by 27% and muscle-specific glucose uptake was increased ∼2.0-fold (Figures 1E, 1F, S1D, and S1E). Muscle glycogen content was increased by 38% in MTKO animals, compared to controls (Figure 1G). The data support the idea that TUG deletion increases cell surface GLUT4 to enhance muscle glucose uptake and whole-body glucose turnover. As in UBX mice, water intake was increased in MTKO mice (Figure S1F), consistent with the prediction that IRAP is also targeted to the plasma membrane and inactivates circulating vasopressin (Habtemichael et al., 2015). We conclude that in muscles lacking TUG, similar to muscles with constitutive TUG cleavage, GSV cargoes are targeted to the cell surface and affect glucose metabolism and physiology.

**Figure 1.**
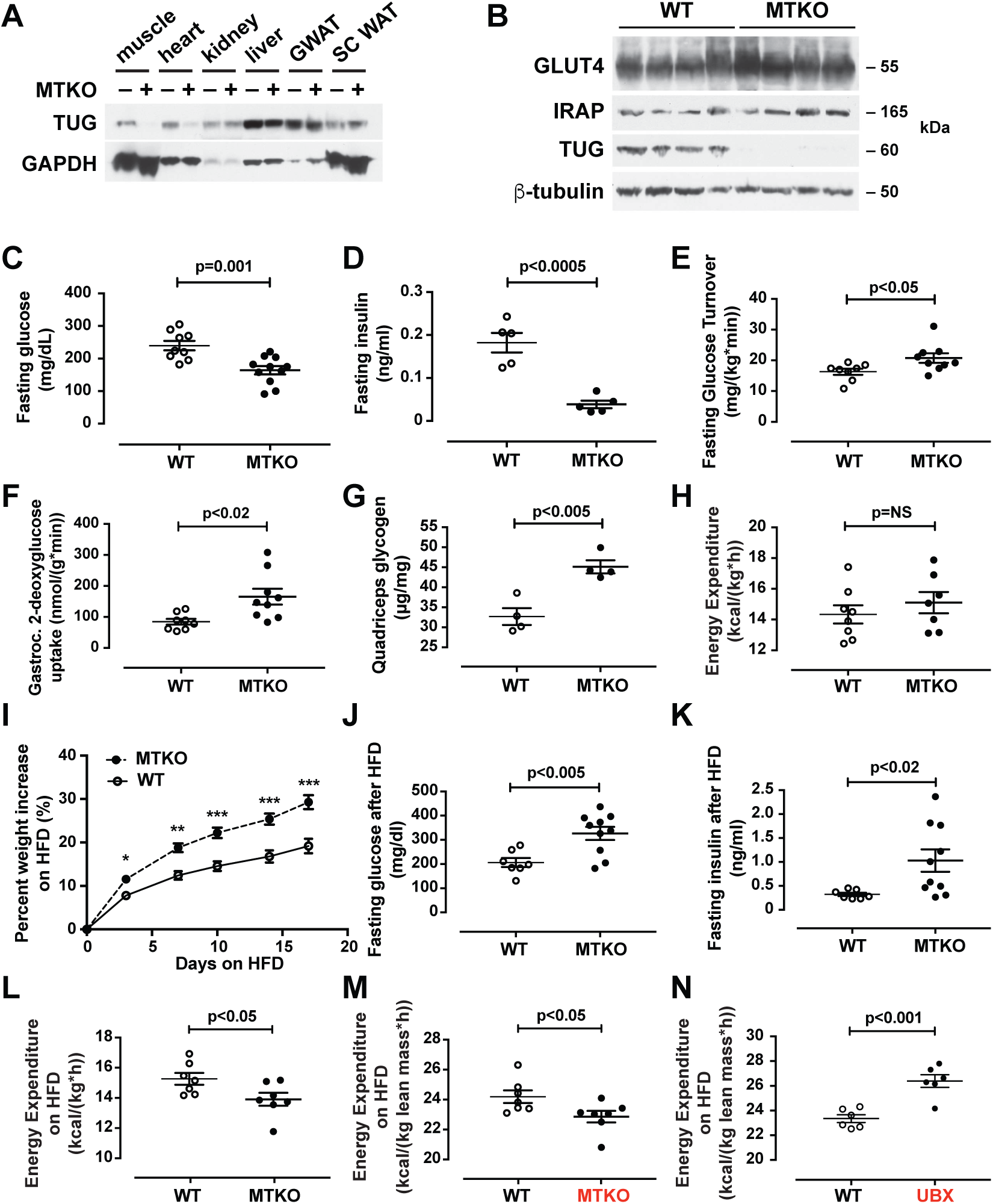
TUG deletion in muscle causes increased glucose uptake and, unlike TUG cleavage, decreased energy expenditure. (A and B) Immunoblots were done as indicated on Muscle TUG Knockout (MTKO) and wildtype (WT) control mice. GWAT and SC WAT indicate gonadal and subcutaneous white adipose tissue, respectively. (C and D) 16-week-old mice were fasted 4-6 h, then plasma glucose and insulin concentrations were measured. (E and F). Tracer infusions were used to measure glucose turnover and gastrocnemius muscle 2-deoxyglucose uptake in fasting 19-week-old mice. (G) Glycogen content was measured in quadriceps muscles of mice fasted for 2 h. (H) Energy expenditure was measured using indirect calorimetry in 17-week old mice. (I) Mice were fed a high-fat diet (HFD) and weight gain was measured. (J and K) Fasting plasma glucose and insulin concentrations were measured after 17 days on a HFD, in 17.5-week old mice. (L and M) Energy expenditure was measured in 14-week-old HFD-fed mice and was normalized to total body weight and lean mass, as indicated. (N) Energy expenditure in 14-week-old HFD-fed UBX mice *(3)* was normalized to lean mass to facilitate comparison. Data are analyzed using t tests and presented as mean ± SEM.

If the increased energy expenditure observed in UBX mice results from cell surface targeting of a GSV cargo, then energy expenditure should be increased in the MTKO mice. This was not observed (Figures 1H and S1G–S1L). At 18 weeks of age, high-fat diet (HFD)-fed MTKO mice gained weight more rapidly than WT controls and developed increased fasting plasma glucose and insulin concentrations (Figures 1I–1K and S1M–S1P). By contrast, HFD-fed UBX mice do not gain excess weight, despite a 14% increase in food intake, and they continue to have reduced fasting plasma glucose concentrations, compared to WT controls (Löffler et al., 2013). Body composition analysis revealed that HFD-fed MTKO mice had increased fat mass (Figures S1Q–S1U). Thus, unlike UBX mice, MTKO have increased susceptibility to diet-induced obesity, compared to WT control animals.

To study energy expenditure, we used younger animals and made measurements prior to the development of significant differences in body weight and composition. In metabolic cages, HFD-fed MTKO mice had a 9% reduction in energy expenditure and no change in respiratory exchange ratio, food intake, or locomotor activity, compared to controls (Figures 1L and S2A–S2H). When normalized to lean mass, energy expenditure was decreased by 6% in MTKO mice and increased by 13% in UBX mice, compared to WT controls (Figures 1M and 1N). In linear regression analyses, we observed these differences in per mouse energy expenditure across a range of body weights (Figure S2I). Intriguingly, on a per mouse basis, HFD-fed MTKO animals had a 22% increase in food intake (Figure S2J). We also observed increased weight gain and gonadal white adipose tissue (GWAT) mass in regular chow (RC) -fed MTKO mice housed under thermoneutral conditions (Figures S2K–S2N). Thus, MTKO and UBX mice have opposite energy expenditure phenotypes, despite similar effects due to cell surface targeting of GSV cargoes, implying that the effect on energy expenditure is not due to a GSV cargo protein.

To learn how gene expression is altered in UBX vs. WT muscles, we analyzed transcriptomes. In UBX muscles, RNAs encoding several proteins for oxidative metabolism and thermogenesis were upregulated (Figure 2A), including sarcolipin (Sln, 3.7-fold increase), Ucp1 (5.7-fold), and Atp2a2 (encoding SERCA2b, 2.6-fold) (Bal et al., 2012; Ikeda et al., 2017; Nedergaard and Cannon, 2014). Pathway analyses identified PPAR signaling as an enriched program of gene expression (Table S1). In transfected cells, TUG accumulates in the nucleus (Orme and Bogan, 2012), and we observed the endogenous TUG C-terminal cleavage product in nuclear fractions of insulin-stimulated muscles (Figures 2B and S3A). The TUG product bound PPARγ and its cofactor, PGC-1α, in transfected cells and using recombinant proteins (Figures S3B–F). Immunoprecipitation of endogenous TUG copurified both PGC-1α and PPARγ from insulin-treated muscles, supporting the physiological relevance of these interactions (Figure 2C). The TUG product bound to peptides corresponding to the N-termini of PPARγ1 and PPARγ2 (Figure S3G). PPARγ2 contains a polymorphism, Pro12Ala, that predicts diabetes risk (Altshuler et al., 2000). We observed increased binding of the TUG product to PPARγ2 peptides containing the protective Ala12 residue, compared to the Pro12 residue (Figures 2D, S3H, and S3I). The data suggest that increased binding of the TUG C-terminal product to PPARγ2 Ala12 may help to recruit PGC-1α and possibly other factors, resulting in enhanced expression of genes for oxidative metabolism and thermogenesis.

**Figure 2.**
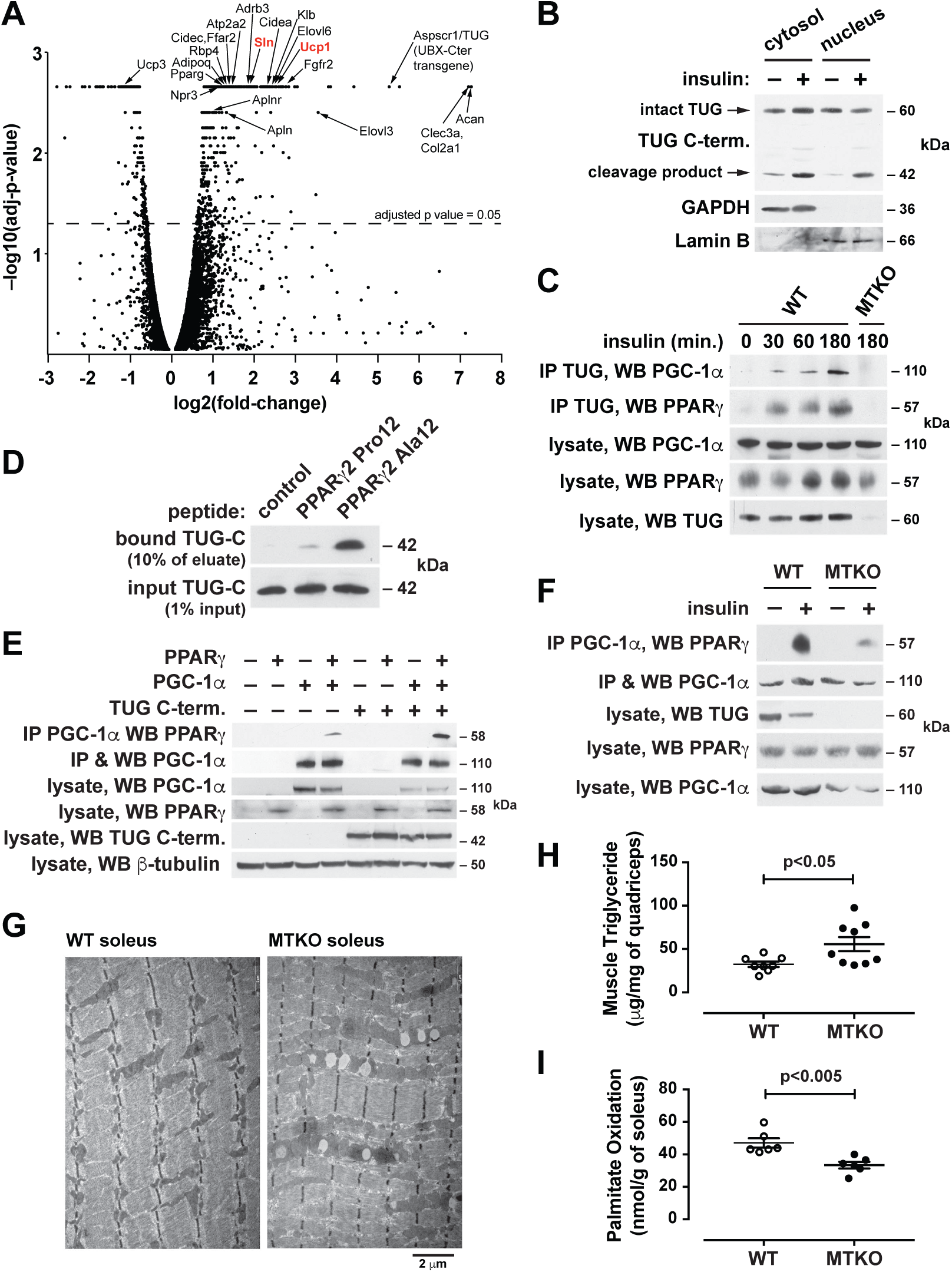
The TUG C-terminal cleavage product acts with PPARγ and PGC-1α to control oxidative metabolism. (A) RNA-seq was used to analyze transcriptomes in quadriceps muscles of UBX and WT mice (n=3 each), and changes in transcript abundance are presented using a volcano plot. Identities of selected transcripts are indicated. (B) Wildtype mice were treated by intraperitoneal (IP) injection of insulin-glucose solution, or saline control, then fractions were isolated from quadriceps muscle and immunoblotted as indicated. (C) Mice were treated by IP injection of insulin-glucose solution, then lysates were prepared from hindlimb muscles at the indicated times. Immunoprecipitation and immunoblots were done as indicated. (D) Peptides corresponding to the N-terminus of PPARγ2, containing Pro12 or Ala12, were immobilized on beads, then incubated with recombinant TUG C-terminal product. Immunoblots were done as indicated. (E) Proteins were expressed by transfection, PGC-1α was immunoprecipitated, and immunoblots were done as indicated. (F) The indicated mice were treated by IP injection of insulin-glucose, or saline control, then euthanized after 3 h. Lysates were prepared from quadriceps, PGC-1α was immunoprecipitated, and immunoblots were done as indicated. (G) The indicated mice were fed a HFD for 2.5 weeks, then soleus muscles were imaged using electron microscopy. (H) Intramyocellular triglyceride was measured in quadriceps from HFD-fed mice. Data are analyzed using t tests and presented as mean ± SEM. (I) Palmitate oxidation was measured *ex vivo* in soleus muscles from mice raised at thermoneutrality.

To test whether the TUG product stabilizes a PPARγ–PGC-1α complex, we immunoprecipitated PGC-1α and immunoblotted PPARγ using transfected cells. Figure 2E shows that a greater number of these complexes were formed when the TUG product was coexpressed. Conversely, fewer PPARγ–PGC-1α complexes were present after insulin stimulation in muscle from MTKO mice, compared to WT controls (Figures 2F and S3J). PGC-1α proteins have limited stability (Adamovich et al., 2013; Sano et al., 2007; Trausch-Azar et al., 2010), and insulin stimulation increased total PGC-1α abundance in muscle in WT, but not MTKO, mice (Figures S3K–N). Consistent with reduced PGC-1α action in MTKO mice, electron microscopy revealed enlarged mitochondria with widened cristae in muscles of HFD-fed MTKO animals, compared to WT controls; lipid droplets were also observed and triglyceride content was increased in muscle lacking TUG (Figures 2G, 2H, and S3O–S3S). *Ex vivo* palmitate oxidation was reduced in soleus muscle of MTKO mice, compared to controls (Figure 2I). The data show that the TUG product binds and stabilizes a PGC-1α–PPARγ complex, increases overall PGC-1α abundance, and promotes fatty acid oxidation.

We next examined whether the TUG C-terminal product regulates the abundance of sarcolipin, which mediates thermogenesis in muscle by uncoupling ATP hydrolysis from Ca^2+^ transport into the sarcoplasmic reticulum (Bal et al., 2018). Insulin stimulated an increase in sarcolipin abundance in WT muscle, which was abrogated by TUG deletion (Figures 3A, 3B, and S4A). Conversely, in UBX muscle, sarcolipin was increased during fasting (Figures 3C and S4B), consistent with RNA-seq data (Figure 2A) and calorimetry (Löffler et al., 2013). After insulin stimulation, the TUG product associated with the sarcolipin promoter (Figure 3D). Insulin also recruited PGC-1α and, to a lesser extent, PPARγ to this site, which was largely independent of TUG (Figures S4C and S4D). In muscle of mice fed a HFD, TUG processing was attenuated and abundance of the TUG protease, Usp25m, was reduced (Figures 3E, 3F, S4E, and S4F). As well, TUG binding at the sarcolipin promoter was reduced (Figure 3G) and the insulin-stimulated increase in sarcolipin protein abundance was attenuated (Figures 3H and 3I). We conclude that the effects on energy expenditure we observe are mediated, in part, by action of the TUG product to increase sarcolipin abundance, and that this action is attenuated when TUG cleavage is impaired in HFD-induced insulin resistance.

**Figure 3.**
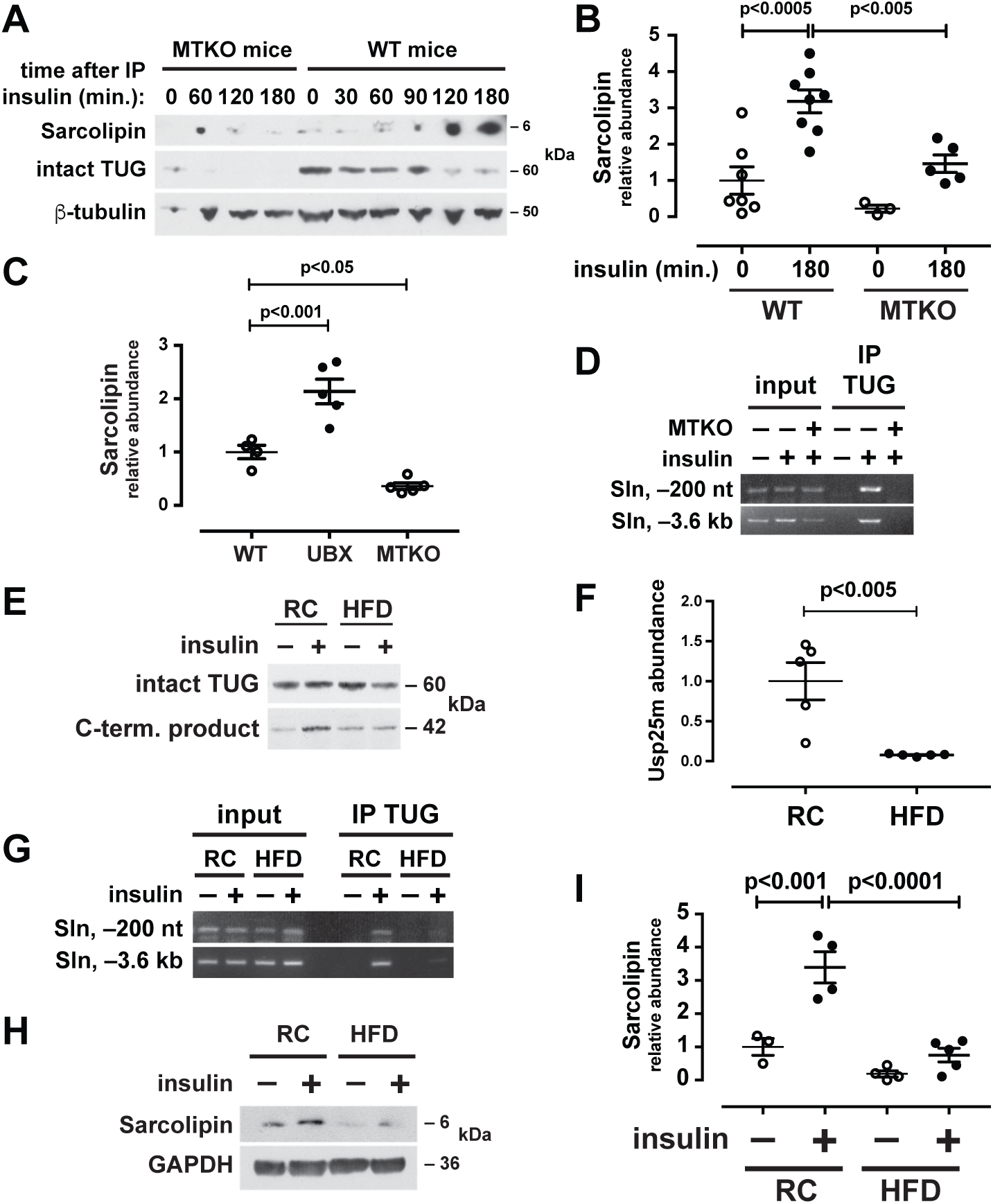
Insulin acts through TUG to enhance production of the thermogenic protein, sarcolipin. (A) Mice were treated with IP insulin-glucose, or saline control, and hindlimb muscles were collected at the indicated times after IP injection. Lysates were immunoblotted as indicated. (B) Studies similar to (A) were repeated in mice raised at 30° C, and sarcolipin abundance was quantified and plotted. Data are analyzed using ANOVA. Mean ± SEM are indicated. (C) The indicated mice were fasted for 4-6 h, then hindlimb muscles were immunoblotted to quantify sarcolipin abundance. Data are analyzed using ANOVA. Mean ± SEM are indicated. (D) The indicated mice were treated with insulin, hindlimb muscles were isolated, and chromatin immunoprecipitation was performed using a TUG C-terminus antibody. PCR detected sequences at the indicated sites relative to the sarcolipin (Sln) transcription start site. (E) Mice fed regular chow (RC) or a high-fat diet (HFD) were treated with IP insulin-glucose, or saline control, then sacrificed after 30 min. Quadriceps lysates were immunoblotted to detect intact TUG and the C-terminal cleavage product. (F) Usp25m abundance was measured by densitometry of immunoblots from replicate samples in (E). Data are plotted relative to RC-fed controls, and are analyzed using a t test. Mean ± SEM are indicated. (G) RC- or HFD-fed mice were fasted, treated with IP insulin-glucose for 30 min. as indicated, then sacrificed. Chromatin immunoprecipitation was done in hindlimb muscles using a TUG C-terminus antibody and PCR to detect sites upstream of the sarcolipin gene. (H) RC- or HFD- fed mice were treated with IP insulin-glucose or saline control, then quadriceps muscles were immunoblotted as indicated. (I) Replicates of data in (H) were quantified, analyzed using ANOVA, and plotted. Mean ± SEM are indicated.

To learn whether the TUG C-terminal product acts by a similar mechanism in adipocytes, we used both primary adipocytes and roscovitine-treated 3T3-L1 cells, which adopt a brown-like phenotype (Wang et al., 2016). In adipocytes, as in muscle, insulin caused entry of the TUG product into the nucleus, where it associated with the Ucp1 promoter (Figures S4G–S4I). Insulin stimulated an increase in Ucp1 protein abundance, which was abrogated by shRNA-mediated depletion of the TUG protease, Usp25m (Figures S4J and S4K). In mice fed a HFD, TUG proteolytic processing was reduced in adipose (Habtemichael et al., 2018), binding of TUG at the Ucp1 promoter was decreased (Figure S4L), and the insulin-stimulated increase in Ucp1 abundance was attenuated (Figures S4M and S4N). The data support the idea that the TUG C-terminal cleavage product acts in adipose, as in muscle, to enhance the expression of thermogenic proteins, and that this effect is reduced because of attenuated TUG cleavage in HFD-induced insulin resistance.

We reasoned that the stability of the TUG C-terminal product and of bound PGC-1α may be controlled by an N-degron pathway (Varshavsky, 2019). Ablation of one such pathway, by acute deletion of the Ate1 arginyltransferase, increases thermogenesis and adipose Ucp1 expression in mice (Brower and Varshavsky, 2009). The TUG product contains an N-terminal serine residue (S165 of intact TUG). Serine is not a known physiologic substrate of Ate1, and we used ubiquitin fusion proteins to test whether the TUG product is subject to this pathway. Deubiquitylase-mediated cleavage of these proteins generates wildtype or mutated TUG products (Figure 4A). In Ate1-deficient murine embryonic fibroblasts (MEFs), compared to control cells, we observed marked accumulation of the wildtype TUG product (Figure 4B). The Ser residue was not fully destabilizing in control MEFs, compared to Asp, a known Ate1 substrate. Yet, accumulation of the wildtype TUG product in Ate1 KO MEFs was similar to that of the S165M mutant, implying that Ate1 controls the main degradative pathway. To further test the effect of Ate1 deletion on TUG C-terminal product stability, we treated cells with cycloheximide. In these “cycloheximide chase” experiments, the stability of the TUG product was greatly enhanced in Ate1 KO MEFs, compared to control cells (Figure 4C). Thus, the stability of the TUG C-terminal product is regulated by an Ate1-dependent degradation pathway.

**Figure 4.**
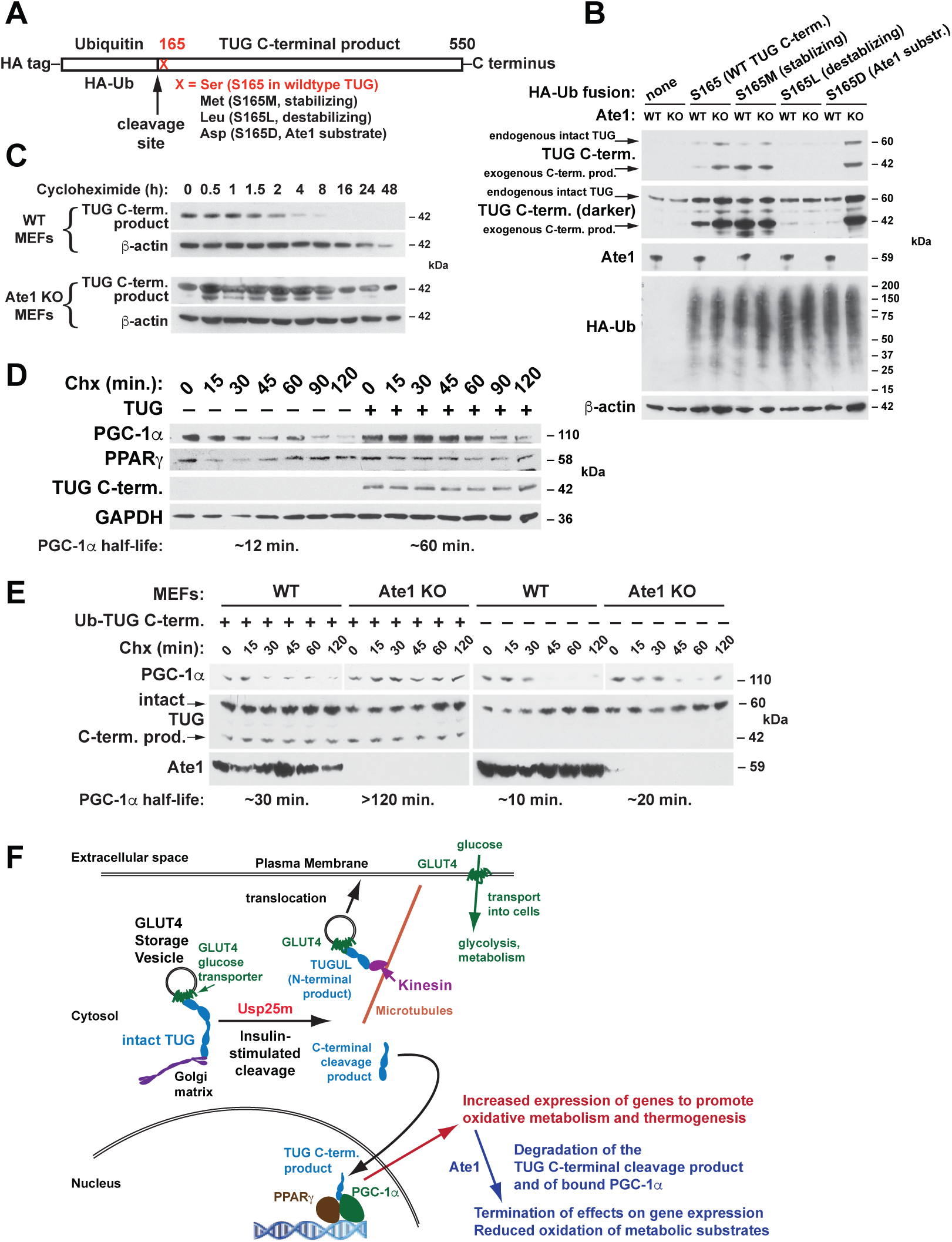
The TUG C-terminal cleavage product stabilizes PGC-1α and is degraded by an Ate1-dependent mechanism. (A) A diagram is shown to indicate how the TUG C-terminal product is produced with different residues at its N-terminus. (B) The indicated constructs were stably expressed in wildtype (WT) and Ate1 knockout (KO) MEFs using retroviruses. Lysates were immunoblotted as indicated. (C) WT and Ate1 KO cells expressing the wildtype TUG C-terminal product were treated with cycloheximide for the times indicated and samples were immunoblotted. (D) HeLa cells were transfected to express PGC-1α and PPARγ, with or without the TUG C-terminal product (beginning with Met), as indicated. Cycloheximide was added for the times indicated and samples were immunoblotted. (E) PGC-1α was expressed, with or without coexpressed HA-Ub-TUG C-term. (WT) fusion protein, in WT and Ate1 KO cells, as indicated. Cycloheximide was added for the times indicated and samples were immunoblotted. (F) An integrated model for how TUG cleavage links glucose uptake and energy expenditure is depicted.

To test whether the half-life of PGC-1α is prolonged by coexpression of the TUG product, we used transfected cells. Cycloheximide chase experiments showed that the half-life of PGC-1α in cells not expressing the TUG product was ∼12 min., similar to previous results (Adamovich et al., 2013; Sano et al., 2007; Trausch-Azar et al., 2010), and this half-life was substantially prolonged by coexpression of the TUG product (Figure 4D). To test whether Ate1 can control PGC-1α stability, we performed similar experiments using WT or Ate1-deficient MEFs. Ate1 deletion had a small effect to stabilize PGC-1α in cells not containing the TUG product, and coexpression of the TUG product both stabilized PGC-1α in WT cells and conferred a marked increase in stability in Ate1 KO cells (Figure 4E). Thus, PGC-1α binds and is stabilized by the TUG C-terminal cleavage product, and the TUG product confers Ate1-dependent stability upon PGC-1α.

## Discussion

Our data support a mechanism whereby glucose uptake and thermogenesis are coordinately regulated by proteolytic cleavage of TUG proteins (Figure 4F). Insulin triggers the cleavage of intact TUG to release GSVs that are trapped at the Golgi, and to promote microtubule-based GLUT4 movement to the plasma membrane (Habtemichael et al., 2018; Löffler et al., 2013). Cleavage generates the TUG C-terminal product, which enters the nucleus, binds and stabilizes a PGC-1α–PPARγ complex, and promotes the expression of genes for oxidative metabolism. Specific genes that are regulated include sarcolipin and Ucp1, in muscle and adipose, respectively. These thermogenic proteins can account, at least in part, for the divergent energy expenditure phenotypes observed in UBX and MTKO mice. The TUG cleavage product has a limited half-life and controls PGC-1α stability, so that both proteins are subject to an Ate1-regulated protein degradation pathway. Thus, the thermogenic effect resulting from the mechanism we describe normally occurs after nutrient intake and has limited duration.

The thermic effect of food is a transient increase in energy expenditure that occurs after meals, which in humans lasts ∼6 h and can account for ∼10% of total energy expenditure (Lowell and Spiegelman, 2000; Reed and Hill, 1996). The mechanism we describe can account, at least in part, for this phenomenon. Insulin signaling upstream of TUG cleavage likely involves a feed-forward circuit, so that cleavage and increased thermogenesis are proportional to glycemic load (Belman et al., 2014; Habtemichael et al., 2018). Attenuated TUG processing may contribute to the reduced thermic effect of food observed in insulin-resistant individuals (de Jonge and Bray, 2002; Du et al., 2014), and to reduced expression of PGC-1α-responsive genes in type 2 diabetes (Mootha et al., 2003). The differential binding of TUG to PPARγ2 Pro12 and Ala12 peptides may explain how this variation can modulate PGC-1α action to control energy expenditure. As the Pro12 variant is suggested to account for ∼25% of type 2 diabetes in the population, understanding the mechanistic basis of this effect is important for public health (Altshuler et al., 2000). Our data also define how Ate1 may be a therapeutic target to modulate the thermic effect of food. We do not know whether Ate1 acts directly on the Ser165 or if modification of this residue is required, as is the case for Cys residues (Hu et al., 2005). As well, we do not know whether the TUG product modifies the transcriptome regulated by PGC-1α–PPARγ complexes, acts with other regulators, or contributes to cross-tissue effects (Bostrom et al., 2012). Nonetheless, our data implicate the TUG C-terminal product as an important regulator of energy expenditure, and imply that understanding this pathway will be significant for metabolic disease.

## Acknowledgments

We thank Michael G. Löffler, Abel Alcázar-Román, Omar Julca-Zevallos, Jacob Culver, Shrikant Mane, Whitney Harris Brown, Eric Li, Roland Calia-Bogan, Anna Kashina, and Derek Toomre for advice and assistance. This work used Core Facilities of the Yale Diabetes Research Center (DRC; NIH P30 DK45735) and services of the CCMI EM facility and Keck Biotechnology Resource at Yale University. This work was supported by NIH R01 DK092661 (to J.S.B.), by American Diabetes Association 1-17-IBS-40 (to J.S.B.), and by R01 DK116774 and R01 DK114793 (to G.I.S.). D.T.L. was supported by T32 GM007205 and F30 DK115037, E.N.H. was supported by a Yale DRC Pilot, under P30 DK45735, and D.F.V. was supported by K23 DK10287. J.-P.C. was supported by São Paulo Research Foundation (FAPESP) grant 2018/04956-5. S.H. was supported by the Brazilian Coordination for the Improvement of Higher Education Personnel (CAPES/PVEX-88881.170862/2018-01) and Postgraduate and Research Dean/Cruzeiro do Sul (PRPGP/UNICSUL-0708/2019).

## Author contributions

E.N.H., D.T.L., and J.S.B. conceptualized the project and designed the experiments. E.N.H., D.T.L., J.-P.C., X.O.W., C.I.S., X.L., F.L.-G., S.G.D., H.L., D.M.R., K.Y.W., B.S.S., S.H., D.F.V., W.P., and J.S.B. performed experiments and analyzed data, G.I.S. supervised metabolic phenotyping experiments, and J.S.B. supervised the overall project and wrote the manuscript with input from all authors.

## Declaration of interests

The authors declare no competing interests.

## Data and materials availability

UBX and TUG fl/fl mice can be provided upon completion of a material transfer agreement to J.S.B. All data are available in the manuscript or the supplementary materials. RNAseq data have been deposited in Gene Expression Omnibus (GEO) under the accession number GSE134846.

## Methods

### Animals

Muscle TUG Knockout (MTKO) mice were produced using a targeting construct obtained from the NIH Knock-out Mouse Program (KOMP; CSD30881). Homologous recombination inserts loxP sites flanking exon 5 of the *Aspscr1* gene, which encodes TUG protein. The construct was electroporated into 129Sv ES cells, recombination was obtained using positive-negative selection, and cells were injected into C57BL/6 blastocysts and implanted into pseudopregnant females. After germline transmission, mice were crossed with Frt transgenic deleter strain to remove the selection cassette. Mice were backcrossed to C57BL/6J for at least 10 generations. To delete TUG in muscle, mice homozygous for floxed TUG allele (TUG^fl/fl^) were bred with TUG^fl/fl^ mice containing a MCK-Cre transgene (Tg(Ckmm-cre)5Khn/J; Jackson stock #006475). Most experiments compared TUG^fl/fl^ (WT) and TUG^fl/fl^ + MCK-Cre (MTKO) mice, and littermates were used as controls. Mice were maintained on at 12 h light/dark cycle and had *ad libitum* access to food and water. Mice were housed at 22° C except for the described experiments done under thermoneutral conditions, for which the mice were housed at 30° C from weaning using Memmert climate chambers. Male mice were used except where indicated. The standard regular chow (RC) diet was Harlan-Teklad 2018S and the high-fat diet (HFD) was Research Diets D12492 (60% kcal from fat). The Yale Institutional Animal Care and Use Committee approved all procedures.

For genotyping, genomic DNA was retrieved by overnight proteinase K digestion of tail biopsies, and was used in PCR to assess the presence of both the floxed TUG allele and the MCK-Cre transgene. For the floxed TUG allele, PCR used 39 cycles with 95° C for 30 seconds, 67° C for 30 seconds, and 72° C for 2.5 minutes, together with the following primer pair: 5’-AGGGCACTGCTCTCATTCTTTG-3’ and 5’-GCCCGCCCAGCTCAGGACAC-3’. For the MCK-Cre transgene, PCR used 38 cycles with 95° C for 30 seconds, 55.5° C for 30 seconds, and 72° C for 2 minutes with the following primer pair: 5’-GCCTTCTCTACACCTGCGG-3’ and 5’-GGTTCGCAAGAACCTGATGG-3’. Genotyping of UBX mice (previously called mTUG^UBX-Cter^ mice) was described previously (Löffler et al., 2013).

### Metabolic and tissue analyses

Fasting blood glucose and insulin concentrations were measured using a handheld glucometer (Onetouch UltraMini, Lifescan) and by an ultrasensitive ELISA (ALPCO, 80-INSMSU-E01). Fat and lean mass were measured using ^1^H NMR (Minispec, Bruker Biospin) (Löffler et al., 2013). Rates of oxygen consumption (VO_2_), carbon dioxide production (VCO_2_), energy expenditure, respiratory exchange ratio, locomotor activity, food consumption, and water intake were measured using CLAMS metabolic cages (Columbus Instruments) (Löffler et al., 2013). Dynamic measurements of glucose flux were done as previously, using a Beckman Glucose Analyzer II (Camporez et al., 2013; Camporez et al., 2017; Löffler et al., 2013). Briefly, a catheter was placed in the right jugular vein 6-7 days before turnover studies were done. After a 6 h fast, 3-[^3^H]glucose (Perkin-Elmer Life Sciences) was infused at a rate of 0.05 μCi/min for 120 min for basal glucose turnover measurement. Next, to measure muscle-specific fasting glucose uptake, 10 μCi of 2-deoxy-D-[1-^14^C] glucose was infused over 20 min., without insulin and with monitoring of plasma glucose and care to minimize any increases. Blood samples were drawn from the tail vein at 5, 15, 25, 35, 45, and 55 min after initiation of 2-deoxyglucose infusion. At the end of the study, mice were treated with intravenous pentobarbital sodium injection (150 mg/kg), tissues were quickly excised, snap frozen in liquid nitrogen, and stored at −80° C for subsequent analysis. Intracellular (6-phosphorylated) 2-deoxyglucose was measured as described previously (Löffler et al., 2013).

Muscle glycogen and triglyceride measurements were done on quadriceps from 4-5 h fasted WT and MTKO mice. For glycogen measurements, chow-fed mice were analyzed using a glycogen assay kit (Biovision, Cat. No. K648-100). For intramyocellular triglyceride, determinations were performed on HFD-fed mice essentially as described (Vatner et al., 2013). Triglycerides were extracted from 80-130 mg quadriceps tissue from each mouse. Tissues were homogenized in ice cold 2:1 chloroform:methanol, and lipids were extracted with shaking at room temperature for 3-4 hours. H_2_SO_4_ was added to ∼100 mM, samples were vortexed, then centrifuged to achieve phase separation, and the organic phase was collected. Aliquots were dried and resuspended in Sekisui Triglyceride-SL reagent (Sekisui) for spectrophotometric determination of triglyceride content. The standard curve was generated using the DC-Cal multi analyte calibrator (Sekisui).

Palmitate oxidation was measured *ex vivo* in soleus muscles from regular chow -fed 20-week old MTKO and WT mice, which had been housed under thermoneutral conditions (30° C) from weaning. Oxidation was measured by collecting released CO_2_ (Chaudry and Gould, 1969; Cuendet et al., 1976). Briefly, soleus muscles were quickly removed and attached to stainless steel clips to maintain resting tension. Muscles were preincubated in 1.5 ml Krebs-Ringer bicarbonate buffer (KRBB) containing 10 mM glucose and 0.5% BSA, pH 7.4, pregassed for 30 min with O_2_, at 35° C. Muscles were then transferred to new vials containing 1.5 ml of the same buffer, but with 0.1 mM palmitic acid and 0.2 μCi/mL [1-^14^C]palmitic acid added. NaOH (0.3 ml at 2 N) was added to an open microtube inside these vials for ^14^CO_2_ adsorption. Incubation was performed for 1 h under the same conditions. At the end of the incubation, muscles were removed, 0.5 ml of 2 N HCl was added to the KRBB, and the incubation was continued for 2 h longer at 35° C. Finally, the NaOH solution (0.3 ml) containing the adsorbed CO_2_ was added to scintillation vials containing scintillation cocktail for radioactivity determination.

For electron microscopy, soleus muscles from HFD-fed WT and MTKO mice, fasted 4-5 h prior to sacrifice, were fixed in 2.5% glutaraldehyde, 2% paraformaldehyde, 0.1 M sodium cacodylate buffer (pH 7.4) at room temperature for 1 h. After rinsing in the same buffer twice, tissue was post-fixed in 1% OsO_4_ at room temperature for 1 h. Specimens were stained *en bloc* using 2% aqueous uranyl acetate for 30 min, dehydrated in a graded series of ethanol to 100%, substituted with propylene oxide, and embedded in EMbed 812 resin. Sample blocks were polymerized in an oven at 60° C overnight. Thin sections (60 nm) were cut using a Leica ultramicrotome (UC7) and post-stained with 2% uranyl acetate and lead citrate. Sections were examined with a FEI Tecnai transmission electron microscope at 80 kV accelerating voltage, and digital images were recorded with an Olympus Morada CCD camera and iTEM imaging software. Image analysis was done blindly using ImageJ/FIJI, and mitochondria were traced manually for measurement of mitochondrial density (number of mitochondria per cross-sectional area, in μm^2^), area per mitochondrion, mitochondrial length (Feret diameter), and mitochondrial width (minimum Feret diameter). Quantification was done for three mice of each genotype, and mitochondria were quantified on 5-9 images (similar to Fig. 2G) from each mouse, so that data from 20 WT and 23 MTKO images were quantified. Averages of the above parameters from each cross-sectional image are plotted in Figures S3P–S3S.

### Reagents and Cell Culture

Antibodies directed to TUG, GLUT4, and IRAP were described previously (Belman et al., 2015; Bogan et al., 2003; Habtemichael et al., 2018; Yu et al., 2007). Other antibodies were purchased, including those directed to GAPDH (Millipore MAB374), β-tubulin (Developmental Studies Hybridoma Bank at Univ. of Iowa, clone E7-b), LaminB1 (Cell Signaling Technology clone D4Q4Z, 12586S), PGC-1α (Life Technologies PA572948), PPARγ (Santa Cruz Biotechnology clone E-8, sc-7273), Sarcolipin (Millipore ABT13), Usp25 (Novus NBP180631 and Abcam ab187156), HA epitope tag (Biolegend clone HA.11, 901502; Cell Signaling clone C29F4, 3724S), ATE1 (Millipore clone 6F11, MABS436), β-actin (Thermo Scientific PIMA515739), flag epitope tag (Cell Signaling clone D6W5B, 14793S and Sigma clone M2, F3165 and A2220), GST (Cell Signaling clone 91G1, 2625S), and insulin receptor β-subunit (Millipore 07-724). Plasmids to express flag-tagged PPARγ, PGC-1α, and PGC-1β were gifts of Dr. Bruce Spiegelman and were obtained from Addgene (plasmids #8895, 1026, 1031, respectively) (Hauser et al., 2000; Lin et al., 2002; Monsalve et al., 2000), as were plasmids for recombinant production of PGC-1α (Addgene #1028 and 1029) (Puigserver et al., 1999). pGEX TUG plasmids were previously described (Orme and Bogan, 2012). The pGEX 4T-1-PPARγ2 plasmid was a gift of Dr. J. Song (Addgene #78773) (Kim et al., 2014). A plasmid encoding HA-tagged ubiquitin was a gift of Dr. Edward Yeh (Addgene #18712) (Kamitani et al., 1997).

3T3-L1, MEF, HEK293, and HeLa cells were cultured in high glucose DMEM GlutaMAX medium (Invitrogen) containing 10% EquaFETAL bioequivalent serum (Atlas Biologicals), antibiotic antimycotic solution (Sigma), and plasmocin (Invivogen). Ate1 knockout (KO) and wildtype (WT) control MEFs were a gift of Dr. Anna Kashina and were described previously (Karakozova et al., 2006). 3T3-L1 adipocytes were differentiated in 10% fetal bovine serum with supplements, as described previously (Belman et al., 2015). For stable expression of exogenous proteins or Usp25 shRNA, MEFs and 3T3-L1 cells were infected with retroviruses and selected using puromycin or FACS (Habtemichael et al., 2018; Liu et al., 2000). Control cells containing empty vector were also subjected to puromycin selection; together with plasmocin, this helped to maintain cells free of mycoplasma. Where indicated, 3T3-L1 adipocytes were differentiated in the presence of 5 μM roscovitine to induce brown adipocyte -like phenotype, as described (Wang et al., 2016). Lipofectamine 2000 (Invitrogen) was used for transient transfection of HEK293 and HeLa cells.

For electroporation, cells were grown to 90% confluence and resuspended in 1 ml of 0.25% trypsin. Cells were pelleted by spinning at 5000 x g and pellet was resuspended in 1 ml of OptiMEM (Gibco). 100 μl of resuspended cells and 2-4 μg of desired plasmid DNA in 5 μL of volume were pipetted into electroporation cuvettes (2 mm gap, Bulldog Bio 12358-346) and electroporated using a NEPA21 electroporation system. Settings used were Poring pulse: V=150, Length=5 ms, Interval=50 ms, Number of pulses=2, Decay rate=10%, Polarity= +. Transfer pulse: V=20, Length=50 ms, Interval=50 ms, Number of pulses=5, Decay rate=40%, Polarity= +/-. After electroporation, cells were diluted with 20 mL of DMEM and plated in 6 well dishes.

### Immunoblots and Immunoprecipitations

For immunoblots of basal and insulin-stimulated tissues, lysates were prepared from mice that had been fasted for 4-6 hours, then treated with intraperitoneal (IP) injection of insulin (8 U/kg) and glucose (1 g/kg) in phosphate buffered saline (PBS), or with an equivalent volume (0.3 ml) of PBS alone, as previously (Löffler et al., 2013). After 30 min (or other indicated durations), mice were anesthetized and sacrificed by cervical dislocation. Glucometer measurements confirmed that no hypoglycemia occurred during the 3 h after IP injections using this protocol. Gonadal white adipose tissue (GWAT), quadriceps, gastrocnemius, and soleus muscles, and other tissues were collected and flash frozen in liquid nitrogen and stored at −80° C.

For immunoblots, tissues were quickly thawed and 200 mg of each tissue were weighed and mixed with lysis buffer (1% IGEPAL CA-630 (Sigma), 20 mM Tris, pH 7.4, 150 mM NaCl, 2 mM EDTA with Complete (Roche) protease inhibitors). A Qiagen TissueLyser II was used to grind the tissue for 3 min. at 30 cycles/sec. To remove insoluble debris, lysates were centrifuged 10 min at 13,000 rpm in a tabletop centrifuge (Eppendorf 5424R) at 4° C. Supernatants were analyzed by SDS-PAGE using Invitrogen NuPAGE gels, transferred to nitrocellulose membranes using a semidry apparatus, and imaged using peroxidase conjugated secondary antibodies and detection on film, as previously (Belman et al., 2015; Bogan et al., 2012; Habtemichael et al., 2018). Quantification was done on exposures within the linear range of the film and used transillumination (Epson Perfection V700 flatbed scanner) together with Silverfast 8 (Lasersoft Imaging) and ImageJ software. Figures were prepared using Adobe Photoshop and Illustrator, and Graphpad Prism software.

For experiments using basal and insulin stimulated 3T3-L1 adipocytes, cells were typically serum starved for 3 h prior to insulin stimulation. Insulin was used at 80-160 nM for 15-30 min unless otherwise specified. Lysis, SDS-PAGE, and immunoblots were done as above.

Immunoprecipitations both tissue and cell lysates were done using the above buffer, and were allowed to proceed overnight at 4° C after addition of the immunoprecipitating antibody. Protein A sepharose (CL-4B, GE LifeSciences) was added and incubations were continued an additional 4 h at 4° C. For immunoprecipitations using epitope tags, affinity matrices were incubated overnight with cell lysates. After pelleting in a benchtop microfuge, beads were washed 6 times with 1% or 0.5% IGEPAL CA-630 buffer and transferred to new tubes. Samples were eluted by heating (5 min., 95° C) in SDS-PAGE sample buffer with 15% 2-mercaptoethanol or without heat using glycine buffer (pH 2.5) with neutralization by Tris base (pH 9). Samples were separated on 4-12% NuPAGE bis-tris gels and immunoblotted as above.

### Fractionations and cycloheximide chase experiments

To fractionate muscle and adipose tissues, 100 mg of tissue was homogenized using 10 strokes in a 2 ml ground glass dounce-type tissue grinder in the buffers provided in the NE-PER nuclear and cytoplasmic extraction kit from ThermoFisher Scientific. Nuclear and cytoplasmic fractions were then prepared according to kit instructions and used for SDS-PAGE and immunoblotting as above. Subcellular fractionation of 3T3-L1 adipocytes was performed as described previously (Belman et al., 2015; Bogan et al., 2001; Yu et al., 2007). Briefly, for each sample, five 10 cm plates of 3T3-L1 adipocytes were homogenized in 5 ml of an ice-cold TES buffer (250 mM sucrose, 10 mM Tris pH 7.4, 0.5 mM EDTA, protease inhibitor cocktail, and 20 mM iodoacetamide) using a glass dounce-type homogenizer. Plasma membrane, light microsome, heavy microsome, nuclear, and cytosolic fractions were isolated by differential centrifugation (Belman et al., 2015; Bogan et al., 2001; Yu et al., 2007). Pellets were resuspended in SDS-PAGE sample buffer with 10% 2-mercaptoethanol. Samples were heated for 5 minutes at 95 °C, separated on 4-12% NuPAGE bis-tris gels, transferred to nitrocellulose membranes, and immunoblotted as above.

For initial tests of protein stability, retroviruses were used to express HA-tagged ubiquitin fused to the TUG C-terminal product (residues 165-550), containing various residues at position 165 as indicated. These fusion constructs were expressed in Ate1 KO MEFs and WT control cells using the pBICD2 vector, and cells with similar levels of expression were selected using FACS of cell surface CD2 (Bogan et al., 2003; Liu et al., 2000; Yu et al., 2007). The relative abundances of HA-tagged ubiquitin were further tested using immunoblots. To assess the rate of TUG C-terminal product degradation, 500 μM cycloheximide was added acutely, cells were lysed at various times after cycloheximide addition, and immunoblots were performed.

To assess effects of TUG and of Ate1 knockout on PGC-1□ stability, cycloheximide chase experiments were using HeLa cells or MEFs. Cells cultured in 10 cm dishes transfected with indicated plasmids using Lipofectamine 2000 or by electroporation, as above. Two days later, cycloheximide was added in 1 ml of prewarmed media to a final concentration of 500 μM. Cells were maintained in cycloheximide for time periods indicated and washed with cold PBS twice prior to lysis, SDS-PAGE, and immunoblotting. Replicate experiments were quantified using densitometry, and protein half-lives were estimated based on a least-squares fit to a first-order exponential decay.

### Transcriptome analysis and chromatin immunoprecipitation

Total RNA was prepared from quadriceps muscles of 4-6 h fasted UBX and WT mice (N=3 each). A NucleoSpin RNA preparation kit (Macherey-Nagel) was used with tissues that had been flash frozen and stored at −80° C. 100 mg of each sample was defrosted on ice and lysed using 10 strokes in a 2 ml glass dounce-type tissue grinder in the buffers provided. For deep sequencing, rRNA were removed using Ribo-Zero (Illumina). Six strand-specific sequencing libraries, 3 replicates per condition, were produced from purified total RNA samples by the Illumina TruSeq stranded protocol. The libraries underwent 76bp single-end sequencing using Illumina HiSeq 2500 according to Illumina protocols, generating between 40-55 million reads per sample. For each read, we trimmed the first 6 nucleotides and the last nucleotides at the point where the Phred score of an examined base fell below 20 using in-house scripts. If, after trimming, the read was shorter than 45-bp, the whole read was discarded. Trimmed reads were mapped to the mouse reference genome (mm10) with a known transcriptome index (UCSC Known Gene annotation) with Tophat v2.1.1 (Trapnell et al., 2009) using the very-sensitive preset, first strand library type, and providing the corresponding gene model annotation. Only the reads that mapped to a single unique location within the genome, with a maximum of two mismatches in the anchor region of the spliced alignment, were reported in these results. We used the default settings for all other Tophat options. Tophat alignments were then processed by Cuffdiff (Cufflinks v2.2.1, Trapnell et al., 2010) to obtain differential gene expression using first strand library type, providing gene model annotation and the genome sequence file for detection and correction of sequence-specific bias that random hexamer can cause during library preparation. Expression between UBX and WT quadriceps was significantly different for 674 transcripts after genome-wide adjustment using Benjamini (to adjusted p<0.05). Of these, 467 had increased and 207 had decreased expression. Pathway analyses were done using DAVID (Huang da et al., 2009), GOrilla (Eden et al., 2009), Ingenuity Pathway Analysis (Qiagen) and MetaCore (Clarivate).

Q-PCR was done as previously (Habtemichael et al., 2018). Briefly, 670 nmol of RNA was used with a high capacity cDNA reverse transcriptase kit (Applied Biosystems). Real time PCR was done using a StepOnePlus system and Power Sybr Green Master Mix (Applied Biosystems). mRNA abundances were normalized to that for β-actin (*Actb*). Primers for PGC-1α (*Ppargc1a*) were AAAAGCTTGACTGGCGTCAT and TCAGGAAGATCTGGGCAAAG and for Actb were TGGAATCCTGTGGCATCCATGAAAC and TAAAACGCAGCTCAGTAACAGTCCG (Liu et al., 2017). In Q-PCR experiments, three technical replicates were done for each biological replicate; technical replicates typically agreed to 3-4 significant figures and were averaged. Biological replicates (from separate mice) are plotted and were used to compare mice.

Chromatin immunoprecipitation was carried out using a ChIP-IT Express Enzymatic Kit (Active Motif). Tissues were quickly thawed and 200 mg of each was weighed, minced, and placed in an Eppendorf tube. Samples were fixed in PBS containing 1.5% formaldehyde for 15 minutes with rotation, then washed three times using PBS. Cleaned fixed tissues were then homogenized using 10-20 strokes in a 2 ml ground glass dounce-type tissue grinder in buffers provided by the ChIP-IT Kit. Precipitations were done using antibodies to the TUG C-terminus, PGC-1α, or PPARγ antibody, together with magnetic beads. Chromatin was washed, eluted, and cross links were reversed per protocol instructions. Eluted DNA as well as pre-IP controls were amplified using GoTaq Green Master Mix (Promega) with primers designed for the promoter regions of Ucp1 and Sarcolipin. Primers used were: Ucp1 −70bp (F – TGTGGCCAGGGCTTTGGGAGT and R – AGATTGCCCGGCACTTCTGCG), Ucp1 - 2.5kb (F – AGCGTCACAGAGGGTCAGT and R – GTGAGGCTGGATCCCCAGA), Ucp1 −5 kb (F – ACATTGCCAAGACTGCGGCCATC and R – ACCCCCAAACAGCAGCAGCAAC), Sarcolipin −200 bp (F – GCCGGAAACAAGAGCTTTCAT and R – TGGGCAGGGCCTAATGTAGT), and Sarcolipin −3.6 kb (F – GCCAGGCCCAAGTGAGAAAGT and R – TGTGGCCAGCAGAGAATAGAGT).

### Recombinant protein expression and pulldowns

GST-tagged constructs were expressed in BL21(DE3)pLys GOLD *E. coli* (Agilent Technologies). 30 ml bacterial starter cultures were grown at 37° C overnight and then added to 1.5 liters of LB media at 37° C. Protein expression was induced with isopropyl-β-D-thiogalactopyranoside at a final concentration of 1 mM once an OD of 0.8 was reached. After 3–4 h, bacteria were lysed in 50 mM Tris, pH 8.0, 300 mM NaCl, 1% Triton X-100 and 1mM PMSF in a French Press. Insoluble debris was pelleted at 16,000 x g for 20 min. GST-tagged protein from supernatant were purified using glutathione-Sepharose 4B (GE Healthcare). Sepharose bound proteins were used as the protein columns. GST-free eluates of protein were prepared by using 1 unit of biotinylated thrombin (Millipore 69672-3) per 100 μl of beads and letting samples incubate on a rotator at 4° C overnight. Thrombin was removed by treating the supernatant with 36 μl of 50% neutravidin bead slurry per 1 unit of thrombin used. Thrombin-cleaved eluted proteins were incubated with protein-bound columns overnight at 4° C, then samples were washed 4-6 times using 0.25% IGEPAL CA-630 buffer. Eluates were analyzed by SDS-PAGE and immunoblotting as above.

For peptide pulldown experiments, biotin-containing synthetic peptides were dissolved to a concentration of 2 mg/mL in Dulbecco’s phosphate buffered saline and 500 μl of peptide solution was incubated with 200 μl of Pierce NeutrAvidin Agarose at 4°C overnight on a rotator. After overnight incubation, beads were washed with 1% IGEPAL CA-630 buffer as above three times, and beads were collected by centrifugation for 1 minute at maximum speed in a benchtop centrifuge. Biotin alone at a concentration of 1 mg/mL was used to generate negative control beads. These pre-incubated beads were then stored at 4°C prior to use. The synthetic peptides used were obtained from LifeTein and were: mmPPARG2, NH2-GETLGDSPVDPEHGAFADALPMSTSQEITMVDTEMPF-Ahx-Ahx-Lys(biotin)-COOH; mmPPARG1, NH2-VDTEMPFWPTNFGISSVDLSVMEDHSHSFDIKPFTTV-Ahx-Ahx-Lys(biotin)-COOH; hsPPARG2P, NH2-GETLGDSPIDPESDSFTDTLSANISQEMT-Ahx-Ahx-Lys(biotin)-COOH; hsPPARG2A, NH2-GETLGDSPIDAESDSFTDTLSANISQEMT-Ahx-Ahx-Lys(biotin)-COOH. Ahx denotes aminohexanoic acid, which was used as a spacer to extend the peptide away from the biotin binding site on the neutravidin beads.

Beads with bound peptides (or biotin only, as a control) were incubated with cell lysates or with recombinant proteins, in different experiments. For cell lysates, 3T3-L1 adipocytes, MEFs, or HEK293 cells were used. Lysates were typically prepared using 0.5% IGEPAL CA-630, although in some experiments this detergent was used at 0.25% or at 1%. For recombinant proteins, 1% IGEPAL CA-630 was used. Whole cell lysates or recombinant proteins were added to 100 μl of beads, prepared as above, and incubated at 4°C overnight on a rotator. Beads were then washed with the same buffer four times, collected by centrifugation in a benchtop centrifuge. Bound proteins were eluted using 100 microliters of sample buffer and heating at 95°C for 5 min. Samples were analyzed by SDS-PAGE and immunoblotting as above.

### Replicates and statistical analysis

All data were replicated in at least two independent experiments, and frequently three or more biological replicates were performed. On scatter plots, data are presented as mean ±SEM. Biological replicates indicate that data were obtained using different mice or different plates of cultured cells. Significance was assessed using an unpaired, two-tailed t-test or using one-way ANOVA with Bonferroni adjustment for multiple comparisons of preselected pairs, and p values indicated in the figures. Differences were considered significant at p<0.05. No statistical tool was used to pre-determine sample sizes; rather, the availability of materials and previous experience determined the number of biological replicates that were used.

## Supplementary Information

**Figure S1.**
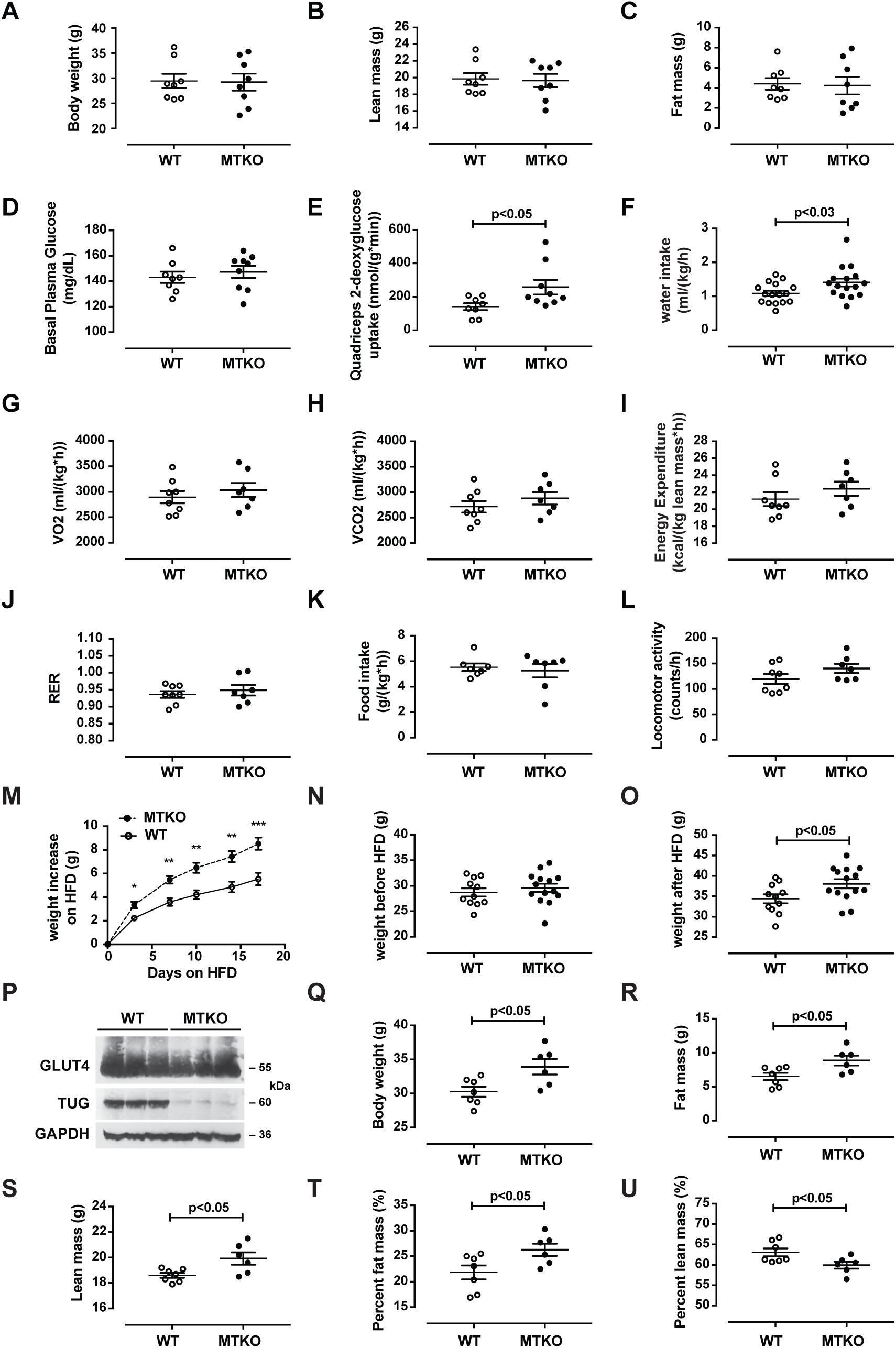
(Related to Figure 1) Metabolic characterization of MTKO mice. (A – C) Body weights and composition were measured in 17-week old WT and MTKO mice. (D) Basal plasma glucose was measured prior to turnover studies in fasting 19-week old mice. (E) Quadriceps-specific 2-deoxyglucose uptake was measured in fasting 19-week old mice. (F) Water intake was measured in 22-week old WT and MTKO mice in metabolic cages. (G – L) The indicated parameters were measured in 17-week old WT and MTKO mice in metabolic cages. (M – O) Weights were measured before and after introduction of a high-fat diet (HFD) in 15-week old MTKO and WT mice. Mice were 18 weeks old at the end of the study. (P) Immunoblots were done as indicated on mice fed a HFD for 3 weeks. (Q – U) Body weights and composition were measured in 18-week old mice that had been fed a HFD for 3 weeks. All data were analyzed using t tests and are presented as mean ± SEM.

**Figure S2.**
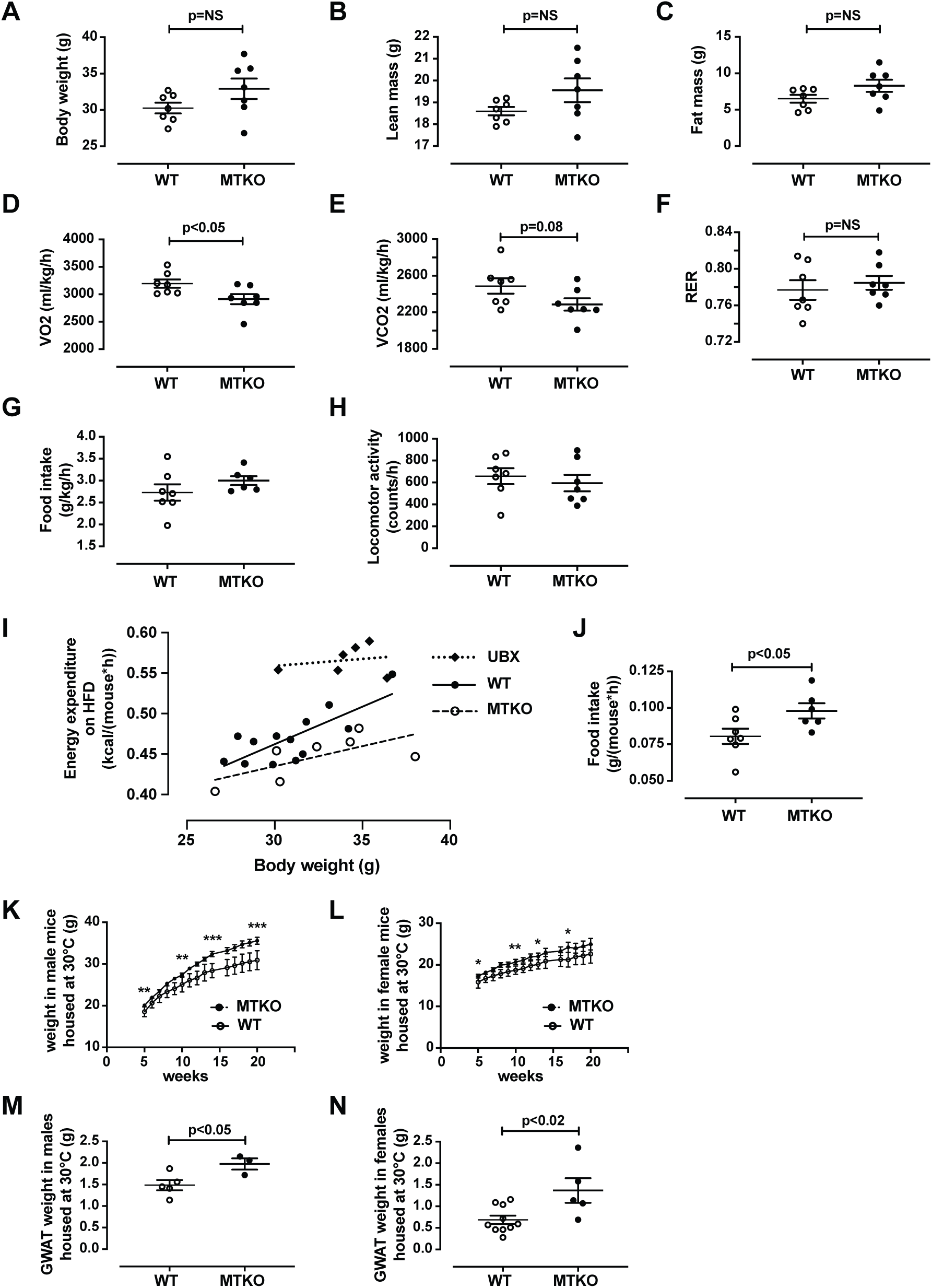
(Related to Figure 1) Characterization of MTKO mice on a high-fat diet or housed at 30° C. (A – C) Body weights and composition of HFD-fed 14-week old MTKO and control (WT) mice. (D – H) HFD-fed 14-week old MTKO and WT mice were housed in metabolic cages and the indicated parameters were measured. (I) Linear regression analysis of energy expenditure in HFD-fed 14-week old UBX, MTKO, and WT mice. (J) Food intake per mouse in 14-week old HFD-fed MTKO and WT mice. (K) Body weights were measured in male MTKO and WT mice housed at 30° C from weaning. All mice were maintained on regular chow. N = 9 MTKO and 11 WT mice. (L) Body weights were measured in female MTKO and WT mice housed at 30° C from weaning. All mice were maintained on regular chow. N = 8 MTKO and 12 WT mice. (M and N) Gonadal white adipose tissues from 20-week old mice used for data in panels K and L were weighed. Data comparing two groups were analyzed using t tests and are presented as mean ± SEM.

**Figure S3.**
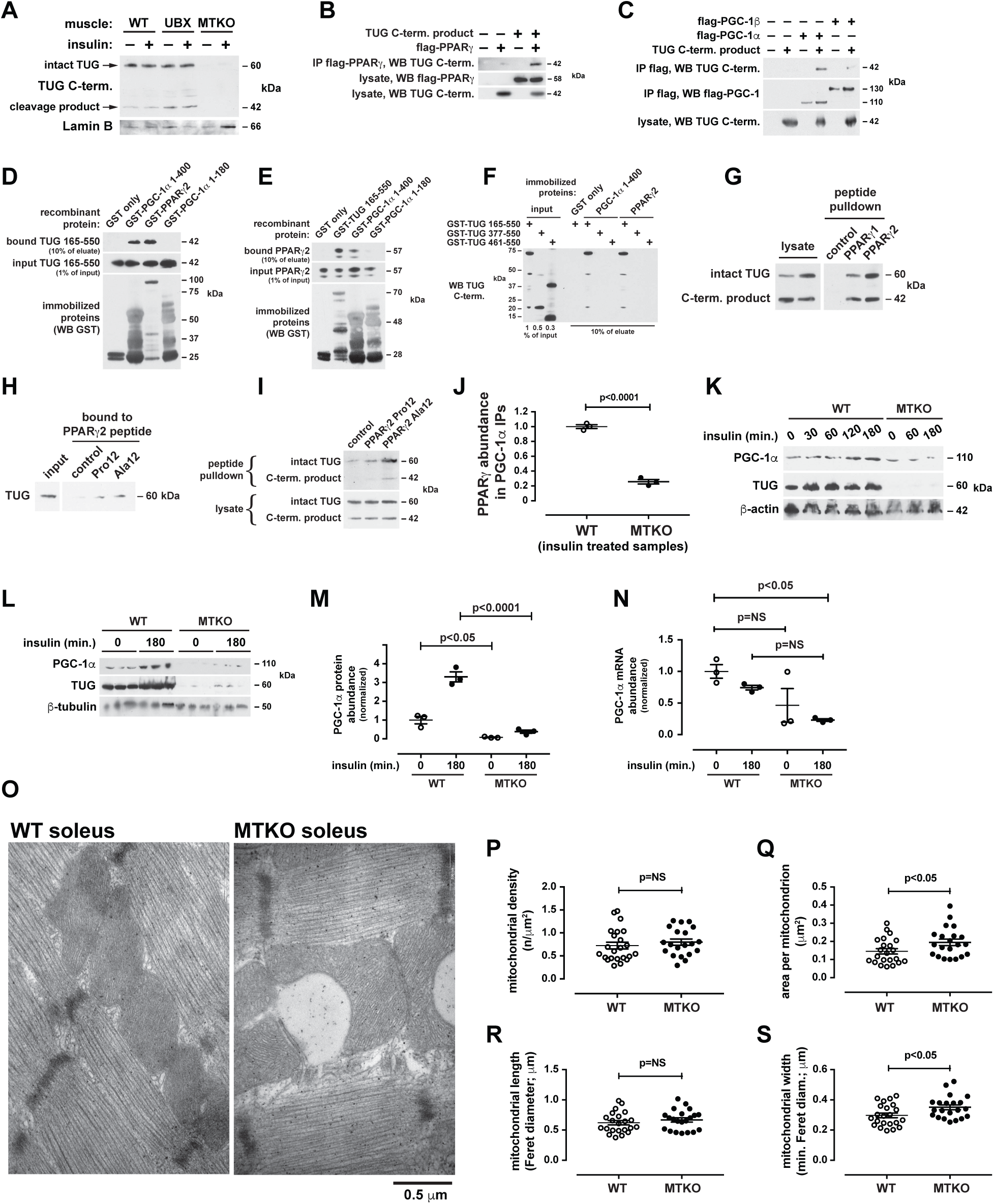
(Related to Figure 2) The TUG C-terminal product binds PPARγ and PGC-1α, and controls PGC-1α abundance and mitochondrial morphology. (A) Nuclear fractions were prepared from quadriceps of mice that had been fasted, treated with intraperitoneal injection of insulin-glucose solution or saline control, and euthanized 30 min. after injection. WT, UBX, and MTKO mice were used, and immunoblots were done as indicated. (B and C) Proteins were expressed by transient transfection of 293 cells, and immunoprecipitations (IP) and western blots (WB) were performed, as indicated. (D) Recombinant proteins were produced as GST fusions, immobilized on glutathione beads, and incubated with a recombinant protein corresponding to the TUG C-terminal cleavage product (residues 165-550). Bound TUG protein was eluted and western blots were performed as indicated. (E) Recombinant proteins were produced, immobilized, and incubated with soluble recombinant PPARγ2 protein. Bound PPARγ2 protein was eluted and western blots were performed as indicated. (F) Truncated forms of TUG were produced as GST fusions and the GST was cleaved off to yield soluble TUG fragments. These were incubated with immobilized GST, PGC-1α, and PPARγ2 as indicated. Bound proteins were eluted and immunoblotted as indicated. (G) Peptides corresponding to the 37 residues at the N-termini of PPARγ1 or PPARγ2, containing a C-terminal biotin, were immobilized on streptavidin beads. Lysates were prepared from MEFs stably expressing the TUG C-terminal product (beginning with a Met residue) using a retrovirus, and were incubated with the beads. Bound proteins were eluted and immunoblotted as indicated. (H) Peptides corresponding to the 29 N-terminal residues of PPARγ2, containing Pro12 or Ala12 residues, were immobilized on beads and incubated with lysates of MEFs stably expressing the TUG C-terminal product. Bound proteins were eluted and immunoblotted as indicated. (I) Peptides used in (H) were incubated with lysates prepared from HEK 293 cells, and bound endogenous (human) intact TUG was eluted and immunoblotted. (J) WT and MTKO mice were treated by IP injection of insulin-glucose solution, then euthanized after 3 h. Lysates were prepared from quadriceps muscles, PGC-1α was immunoprecipitated, and immunoblots were done to detect PPARγ, as shown in Fig. 2F. The relative abundances of PPARγ in replicate experiments were quantified using densitometry and are plotted. (K) WT and MTKO mice were treated by IP injection of insulin-glucose, or saline control, then sacrificed at the indicated times after injection. Quadriceps muscles were immunoblotted as indicated. (L and M) WT and MTKO mice were treated by IP injection of insulin-glucose, or saline control, then sacrificed after 3 h. Lysates were prepared from quadriceps muscles, PGC-1α was immunoblotted, and the relative abundances in each sample were quantified using densitometry. (N) WT and MTKO mice were treated with IP insulin-glucose, or saline control, then sacrificed 3 h later. RNA was prepared from quadriceps muscles, and Q-PCR was used to measure PGC-1α (*Ppargc1a*) mRNA abundance. (O) WT and MTKO soleus muscles from mice that had been fed a HFD for 2.5 weeks were imaged using electron microscopy. Lipid droplets were noted in MTKO muscles, but not WT muscles, and were adjacent to mitochondria, as shown. (P – S) Images of soleus muscles from HFD-fed WT and MTKO mice (N=3 each) were obtained by electron microscopy and were analyzed to quantify mitochondrial density, area, length, and width. Mitochondria were traced manually on 5-9 images from each mouse. Each data point represents the average of the measurements from a single image. Data were analyzed using t tests or ANOVA with adjustment for multiple comparisons, as appropriate, and are presented as mean ± SEM.

**Figure S4.**
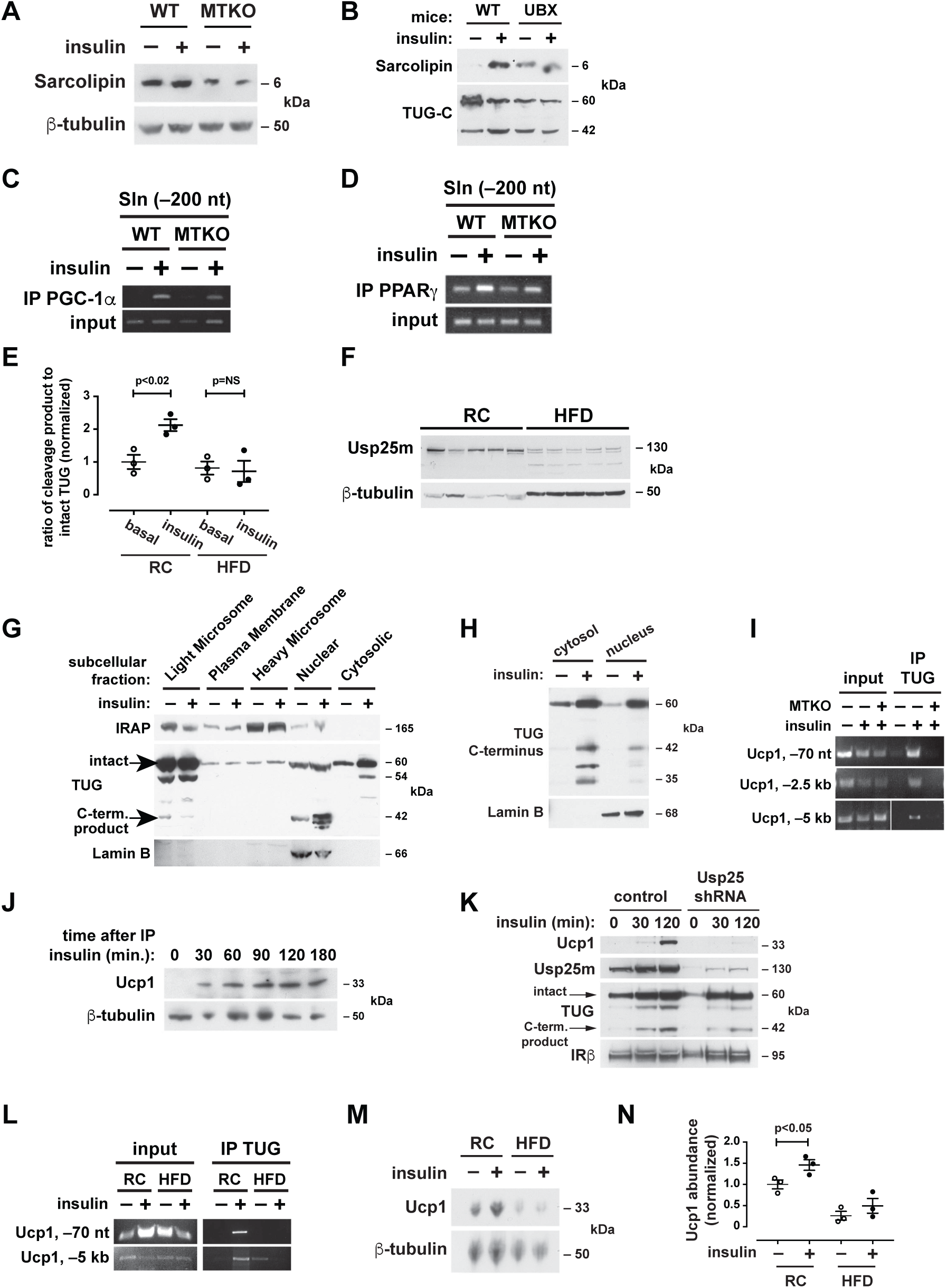
(Related to Figure 3) Sarcolipin and Ucp1 abundances are regulated by TUG and are decreased in diet-induced insulin resistance. (A) WT and MTKO mice were housed at 30° C from the time of weaning. At age 20 weeks, mice were fasted, treated with IP insulin-glucose or saline control, and sacrificed 3 h later. Hindlimb muscle lysates were immunoblotted as indicated. (B) WT and UBX mice were treated with IP insulin-glucose, or saline control, sacrificed after 30 min., and quadriceps muscles were immunoblotted as indicated. (C and D) WT and MTKO mice were treated with IP insulin-glucose, or saline control, and sacrificed after 30 min. Hindlimb muscles were used for chromatin immunoprecipitation, together with PGC-1α and PPARγ antibodies, as indicated. PCR was used to detect an amplicon at −200 nucleotides relative to the sarcolipin transcription start site. (E) WT mice were fed regular chow (RC) or a high-fat diet (HFD) for 3 weeks, then treated for 30 min with IP insulin-glucose, or saline control. Quadriceps muscles were immunoblotted to detect intact TUG and the 42 kDa C-terminal cleavage product, and bands were quantified using densitometry. The ratio of cleavage product to intact TUG is plotted, normalized to basal control samples. (F) WT mice were fed RC or a HFD for 3 weeks, fasted, and sacrificed. Hindlimb muscles were isolated and immunoblotted as indicated. (G) Subcellular fractions were prepared from basal and insulin-treated 3T3-L1 adipocytes, and were immunoblotted as indicated to detect the TUG C-terminus. (H) Mice were fasted, treated by IP injection of insulin-glucose solution or saline control for 30 min., then gonadal white adipose tissue was isolated. Cytosolic and nuclear fractions were prepared and immunoblotted as indicted. (I) WT and MTKO mice were fasted, treated by IP injection of insulin-glucose solution or saline control for 30 min., then hindlimb muscles were isolated. Chromatin immunoprecipitations were performed using an antibody to the TUG C-terminus, and PCR was used to detect sequences at the indicated locations upstream of the Ucp1 transcription start site. (J) Mice were treated with IP injection of insulin-glucose for the indicated time periods and gonadal white adipose tissue was immunoblotted as shown. (K) 3T3-L1 adipocytes were differentiated in the presence of roscovitine to induce a brown-like phenotype, stimulated with insulin for the indicated times, and immunoblotted as shown. (L) Chromatin immunoprecipitation was done as in (I) using GWAT from mice maintained on regular chow (RC) or fed a high-fat diet (HFD) for 3 weeks. Mice were treated with insulin-glucose or saline for 30 min., as indicated prior to sample preparation. (M) Immunoblots were done on gonadal white adipose tissue from RC or HFD-fed mice, treated prior to euthanasia with IP injection of saline or glucose-insulin solution, as indicated. (N) Quantification of relative Ucp1 protein abundances in replicates of the experiment shown in (M).

**Table S1.**
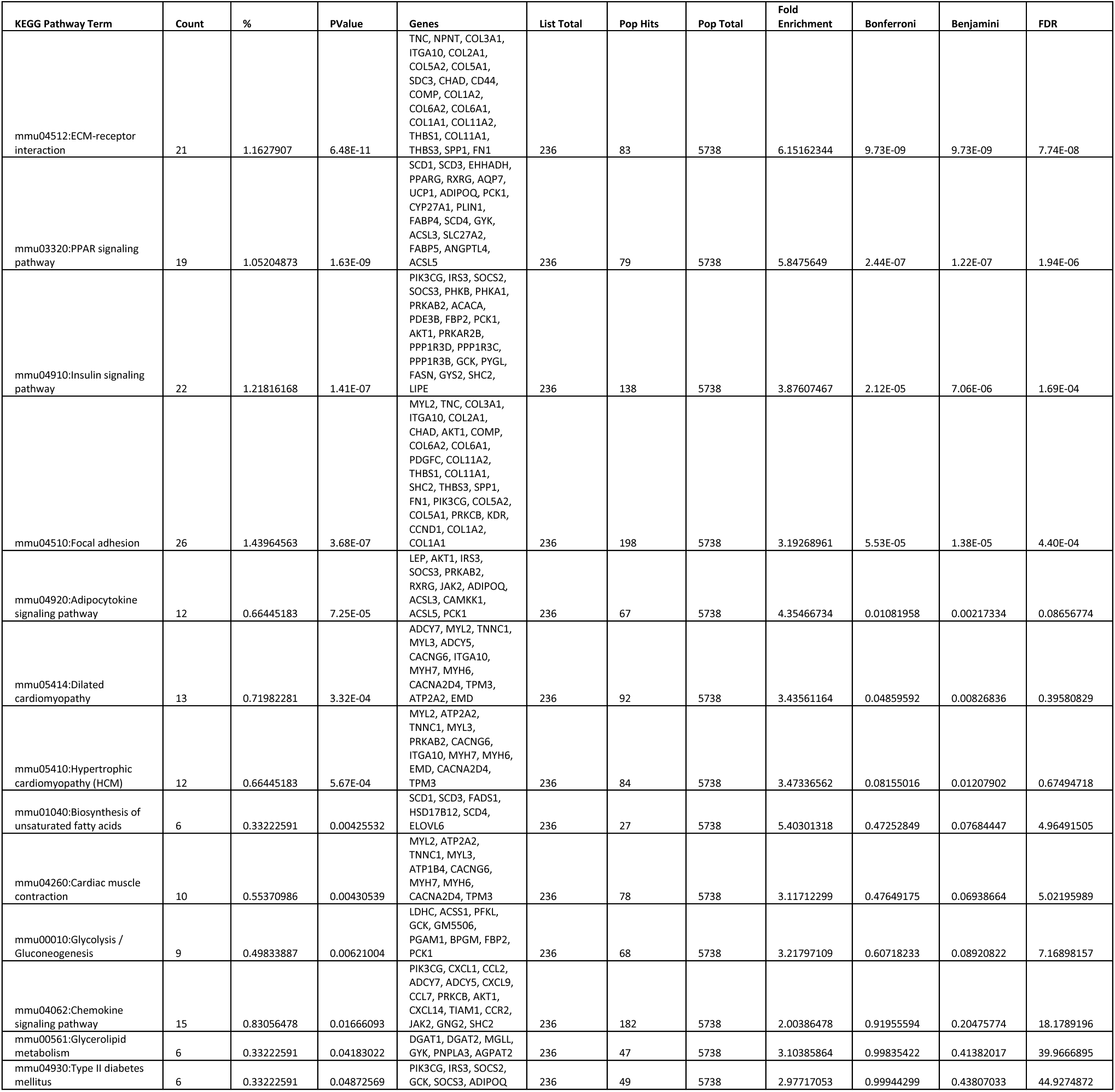
(Related to Figure 2) Pathway analysis of transcriptome alterations in quadriceps of UBX mice, compared to WT controls. Genes with significantly different transcript abundances in UBX and WT mice were identified using RNA-seq (N=3 samples in each group), and were mapped using DAVID and the KEGG database.

## Notes

#### Summary of Updates

Corrected an error in the abstract.

## References

Adamovich, Y., Shlomai, A., Tsvetkov, P., Umansky, K.B., Reuven, N., Estall, J.L., Spiegelman, B.M., and Shaul, Y. (2013). The protein level of PGC-1alpha, a key metabolic regulator, is controlled by NADH-NQO1. Mol Cell Biol 33, 2603–2613.

Altshuler, D., Hirschhorn, J.N., Klannemark, M., Lindgren, C.M., Vohl, M.C., Nemesh, J., Lane, C.R., Schaffner, S.F., Bolk, S., Brewer, C., et al. (2000). The common PPARgamma Pro12Ala polymorphism is associated with decreased risk of type 2 diabetes. Nat Genet 26, 76–80.

Bal, N.C., Maurya, S.K., Sopariwala, D.H., Sahoo, S.K., Gupta, S.C., Shaikh, S.A., Pant, M., Rowland, L.A., Bombardier, E., Goonasekera, S.A., et al. (2012). Sarcolipin is a newly identified regulator of muscle-based thermogenesis in mammals. Nat Med 18, 1575–1579.

Bal, N.C., Sahoo, S.K., Maurya, S.K., and Periasamy, M. (2018). The Role of Sarcolipin in Muscle Non-shivering Thermogenesis. Front Physiol 9, 1217.

Belman, J.P., Bian, R.R., Habtemichael, E.N., Li, D.T., Jurczak, M.J., Alcazar-Roman, A., McNally, L.J., Shulman, G.I., and Bogan, J.S. (2015). Acetylation of TUG protein promotes the accumulation of GLUT4 glucose transporters in an insulin-responsive intracellular compartment. J Biol Chem 290, 4447–4463.

Belman, J.P., Habtemichael, E.N., and Bogan, J.S. (2014). A proteolytic pathway that controls glucose uptake in fat and muscle. Reviews in endocrine & metabolic disorders 15, 55–66.

Bogan, J.S. (2012). Regulation of glucose transporter translocation in health and diabetes. Annu Rev Biochem 81, 507–532.

Bogan, J.S., Hendon, N., McKee, A.E., Tsao, T.S., and Lodish, H.F. (2003). Functional cloning of TUG as a regulator of GLUT4 glucose transporter trafficking. Nature 425, 727–733.

Bogan, J.S., McKee, A.E., and Lodish, H.F. (2001). Insulin-responsive compartments containing GLUT4 in 3T3-L1 and CHO cells: regulation by amino acid concentrations. Mol Cell Biol 21, 4785–4806.

Bogan, J.S., Rubin, B.R., Yu, C., Loffler, M.G., Orme, C.M., Belman, J.P., McNally, L.J., Hao, M., and Cresswell, J.A. (2012). Endoproteolytic cleavage of TUG protein regulates GLUT4 glucose transporter translocation. J Biol Chem 287, 23932–23947.

Bostrom, P., Wu, J., Jedrychowski, M.P., Korde, A., Ye, L., Lo, J.C., Rasbach, K.A., Bostrom, E.A., Choi, J.H., Long, J.Z., et al. (2012). A PGC1-alpha-dependent myokine that drives brown-fat-like development of white fat and thermogenesis. Nature 481, 463–468.

Brower, C.S., and Varshavsky, A. (2009). Ablation of arginylation in the mouse N-end rule pathway: loss of fat, higher metabolic rate, damaged spermatogenesis, and neurological perturbations. PLoS One 4, e7757.

Camporez, J.P., Jornayvaz, F.R., Lee, H.Y., Kanda, S., Guigni, B.A., Kahn, M., Samuel, V.T., Carvalho, C.R., Petersen, K.F., Jurczak, M.J., et al. (2013). Cellular mechanism by which estradiol protects female ovariectomized mice from high-fat diet-induced hepatic and muscle insulin resistance. Endocrinology 154, 1021–1028.

Camporez, J.P., Wang, Y., Faarkrog, K., Chukijrungroat, N., Petersen, K.F., and Shulman, G.I. (2017). Mechanism by which arylamine N-acetyltransferase 1 ablation causes insulin resistance in mice. Proc Natl Acad Sci U S A 114, E11285–E11292.

Chaudry, I.H., and Gould, M.K. (1969). Kinetics of glucose uptake in isolated soleus muscle. Biochim Biophys Acta 177, 527–536.

Cuendet, G.S., Loten, E.G., Jeanrenaud, B., and Renold, A.E. (1976). Decreased basal, noninsulin-stimulated glucose uptake and metabolism by skeletal soleus muscle isolated from obese-hyperglycemic (ob/ob) mice. J Clin Invest 58, 1078–1088.

de Jonge, L., and Bray, G.A. (2002). The thermic effect of food is reduced in obesity. Nutr Rev 60, 295–297; author reply 299-300.

Du, S., Rajjo, T., Santosa, S., and Jensen, M.D. (2014). The thermic effect of food is reduced in older adults. Horm Metab Res 46, 365–369.

Eden, E., Navon, R., Steinfeld, I., Lipson, D., and Yakhini, Z. (2009). GOrilla: a tool for discovery and visualization of enriched GO terms in ranked gene lists. BMC Bioinformatics 10, 48.

Habtemichael, E.N., Alcazar-Roman, A., Rubin, B.R., Grossi, L.R., Belman, J.P., Julca, O., Loffler, M.G., Li, H., Chi, N.W., Samuel, V.T., et al. (2015). Coordinated Regulation of Vasopressin Inactivation and Glucose Uptake by Action of TUG Protein in Muscle. J Biol Chem 290, 14454–14461.

Habtemichael, E.N., Li, D.T., Alcazar-Roman, A., Westergaard, X.O., Li, M., Petersen, M.C., Li, H., DeVries, S.G., Li, E., Julca-Zevallos, O., et al. (2018). Usp25m protease regulates ubiquitin-like processing of TUG proteins to control GLUT4 glucose transporter translocation in adipocytes. J Biol Chem 293, 10466–10486.

Hauser, S., Adelmant, G., Sarraf, P., Wright, H.M., Mueller, E., and Spiegelman, B.M. (2000). Degradation of the peroxisome proliferator-activated receptor gamma is linked to ligand-dependent activation. J Biol Chem 275, 18527–18533.

Hu, R.G., Sheng, J., Qi, X., Xu, Z., Takahashi, T.T., and Varshavsky, A. (2005). The N-end rule pathway as a nitric oxide sensor controlling the levels of multiple regulators. Nature 437, 981–986.

Huang da, W., Sherman, B.T., and Lempicki, R.A. (2009). Systematic and integrative analysis of large gene lists using DAVID bioinformatics resources. Nature protocols 4, 44–57.

Ikeda, K., Kang, Q., Yoneshiro, T., Camporez, J.P., Maki, H., Homma, M., Shinoda, K., Chen, Y., Lu, X., Maretich, P., et al. (2017). UCP1-independent signaling involving SERCA2b-mediated calcium cycling regulates beige fat thermogenesis and systemic glucose homeostasis. Nat Med 23, 1454–1465.

Kamitani, T., Kito, K., Nguyen, H.P., and Yeh, E.T. (1997). Characterization of NEDD8, a developmentally down-regulated ubiquitin-like protein. J Biol Chem 272, 28557–28562.

Karakozova, M., Kozak, M., Wong, C.C., Bailey, A.O., Yates, J.R., 3rd, Mogilner, A., Zebroski, H., and Kashina, A. (2006). Arginylation of beta-actin regulates actin cytoskeleton and cell motility. Science 313, 192-196.

Kim, J.H., Park, K.W., Lee, E.W., Jang, W.S., Seo, J., Shin, S., Hwang, K.A., and Song, J. (2014). Suppression of PPARgamma through MKRN1-mediated ubiquitination and degradation prevents adipocyte differentiation. Cell Death Differ 21, 594–603.

Leto, D., and Saltiel, A.R. (2012). Regulation of glucose transport by insulin: traffic control of GLUT4. Nat Rev Mol Cell Biol 13, 383–396.

Li, D.T., Habtemichael, E.N., and Bogan, J.S. (2019). Vasopressin inactivation: Role of insulin-regulated aminopeptidase. Vitam Horm 113, in press, doi: 10.1016/bs.vh.2019.1008.1017.

Lin, J., Puigserver, P., Donovan, J., Tarr, P., and Spiegelman, B.M. (2002). Peroxisome proliferator-activated receptor gamma coactivator 1beta (PGC-1beta), a novel PGC-1-related transcription coactivator associated with host cell factor. J Biol Chem 277, 1645–1648.

Liu, S., Kim, T.H., Franklin, D.A., and Zhang, Y. (2017). Protection against High-Fat-Diet-Induced Obesity in MDM2(C305F) Mice Due to Reduced p53 Activity and Enhanced Energy Expenditure. Cell reports 18, 1005–1018.

Liu, X., Constantinescu, S.N., Sun, Y., Bogan, J.S., Hirsch, D., Weinberg, R.A., and Lodish, H.F. (2000). Generation of mammalian cells stably expressing multiple genes at predetermined levels. Anal Biochem 280, 20–28.

Löffler, M.G., Birkenfeld, A.L., Philbrick, K.M., Belman, J.P., Habtemichael, E.N., Booth, C.J., Castorena, C.M., Choi, C.S., Jornayvaz, F.R., Gassaway, B.M., et al. (2013). Enhanced fasting glucose turnover in mice with disrupted action of TUG protein in skeletal muscle. J Biol Chem 288, 20135–20150.

Lowell, B.B., and Spiegelman, B.M. (2000). Towards a molecular understanding of adaptive thermogenesis. Nature 404, 652–660.

Monsalve, M., Wu, Z., Adelmant, G., Puigserver, P., Fan, M., and Spiegelman, B.M. (2000). Direct coupling of transcription and mRNA processing through the thermogenic coactivator PGC-1. Mol Cell 6, 307–316.

Mootha, V.K., Lindgren, C.M., Eriksson, K.F., Subramanian, A., Sihag, S., Lehar, J., Puigserver, P., Carlsson, E., Ridderstrale, M., Laurila, E., et al. (2003). PGC-1alpha-responsive genes involved in oxidative phosphorylation are coordinately downregulated in human diabetes. Nat Genet 34, 267–273.

Nedergaard, J., and Cannon, B. (2014). The browning of white adipose tissue: some burning issues. Cell Metab 20, 396–407.

Orme, C.M., and Bogan, J.S. (2012). The ubiquitin regulatory X (UBX) domain-containing protein TUG regulates the p97 ATPase and resides at the endoplasmic reticulum-golgi intermediate compartment. J Biol Chem 287, 6679–6692.

Puigserver, P., Adelmant, G., Wu, Z., Fan, M., Xu, J., O’Malley, B., and Spiegelman, B.M. (1999). Activation of PPARgamma coactivator-1 through transcription factor docking. Science 286, 1368–1371.

Reed, G.W., and Hill, J.O. (1996). Measuring the thermic effect of food. Am J Clin Nutr 63, 164–169.

Sano, M., Tokudome, S., Shimizu, N., Yoshikawa, N., Ogawa, C., Shirakawa, K., Endo, J., Katayama, T., Yuasa, S., Ieda, M., et al. (2007). Intramolecular control of protein stability, subnuclear compartmentalization, and coactivator function of peroxisome proliferator-activated receptor gamma coactivator 1alpha. J Biol Chem 282, 25970–25980.

Trapnell, C., Pachter, L., and Salzberg, S.L. (2009). TopHat: discovering splice junctions with RNA-Seq. Bioinformatics 25, 1105–1111.

Trapnell, C., Williams, B.A., Pertea, G., Mortazavi, A., Kwan, G., van Baren, M.J., Salzberg, S.L., Wold, B.J., and Pachter, L. (2010). Transcript assembly and quantification by RNA-Seq reveals unannotated transcripts and isoform switching during cell differentiation. Nat Biotechnol 28, 511–515.

Trausch-Azar, J., Leone, T.C., Kelly, D.P., and Schwartz, A.L. (2010). Ubiquitin proteasome-dependent degradation of the transcriptional coactivator PGC-1{alpha} via the N-terminal pathway. J Biol Chem 285, 40192–40200.

Varshavsky, A. (2019). N-degron and C-degron pathways of protein degradation. Proc Natl Acad Sci U S A 116, 358–366.

Vatner, D.F., Weismann, D., Beddow, S.A., Kumashiro, N., Erion, D.M., Liao, X.H., Grover, G.J., Webb, P., Phillips, K.J., Weiss, R.E., et al. (2013). Thyroid hormone receptor-beta agonists prevent hepatic steatosis in fat-fed rats but impair insulin sensitivity via discrete pathways. Am J Physiol Endocrinol Metab 305, E89–100.

Wang, H., Liu, L., Lin, J.Z., Aprahamian, T.R., and Farmer, S.R. (2016). Browning of White Adipose Tissue with Roscovitine Induces a Distinct Population of UCP1(+) Adipocytes. Cell Metab 24, 835–847.

Yu, C., Cresswell, J., Loffler, M.G., and Bogan, J.S. (2007). The glucose transporter 4-regulating protein TUG is essential for highly insulin-responsive glucose uptake in 3T3-L1 adipocytes. J Biol Chem 282, 7710–7722.

